# Brain Glutathione and GABA+ levels in autistic children

**DOI:** 10.1101/2023.09.28.559718

**Authors:** Yulu Song, Kathleen E. Hupfeld, Christopher W. Davies-Jenkins, Helge J. Zöllner, Saipavitra Murali-Manohar, Mumuni Abdul-Nashirudeen, Deana Crocetti, Vivek Yedavalli, Georg Oeltzschner, Natalie Alessi, Mitchell A. Batschelett, Nicolaas A.J. Puts, Stewart H. Mostofsky, Richard A.E. Edden

## Abstract

Autism spectrum disorder (ASD) is a neurodevelopmental condition characterized by social communication challenges and repetitive behaviors. Altered neurometabolite levels, including glutathione (GSH) and gamma-aminobutyric acid (GABA), have been proposed as potential contributors to the biology underlying ASD. This study investigated whether cerebral GSH or GABA levels differ between a large cohort of children aged 8-12 years with ASD (n=52) and typically developing children (TDC, n=49). A comprehensive analysis of GSH and GABA levels in multiple brain regions, including the primary motor cortex (SM1), thalamus (Thal), medial prefrontal cortex (mPFC), and supplementary motor area (SMA), was conducted using single-voxel HERMES MR spectroscopy at 3T. The results revealed no significant differences in cerebral GSH or GABA levels between the ASD and TDC groups across all examined regions. These findings suggest that the concentrations of GSH (an important antioxidant and neuromodulator) and GABA (a major inhibitory neurotransmitter) do not exhibit marked alterations in children with ASD compared to TDC. A statistically significant positive correlation was observed between GABA levels in the SM1 and Thal regions with ADHD inattention scores. No significant correlation was found between metabolite levels and hyper/impulsive scores of ADHD, measures of core ASD symptoms (ADOS-2, SRS-P) or adaptive behavior (ABAS-2). While both GSH and GABA have been implicated in various neurological disorders, the current study provides valuable insights into the specific context of ASD and highlights the need for further research to explore other neurochemical alterations that may contribute to the pathophysiology of this complex disorder.

**Lay summary:** Autism spectrum disorder (ASD) is a neurodevelopmental condition characterized by social communication challenges and repetitive behaviors. Altered glutathione (GSH, an important antioxidant and neuromodulator) and gamma-aminobutyric acid (GABA, a major inhibitory neurotransmitter) levels have been proposed as potential contributors to the biology underlying ASD. Here, we used advanced edited Magnetic Resonance Spectroscopy (MRS) to measure levels of these low-concentration metabolites in four brain regions of a pediatric cohort. Contrary to our hypothesis, no significant difference was found between ASD and control subjects in either GSH or GABA levels in any brain region.

## 1. Introduction

Autism spectrum disorder (ASD), or autism, is a lifetime neurodevelopmental disorder characterized by differences in communication and social interaction, along with restrictive and repetitive motor behaviors and atypical sensory experiences(Guze, 1995; Lord et al., 2018). Note that we recognize autism, or Autism Spectrum Condition (ASC) is often the preferred terminology. However, we consider ASD a necessary term given the description in the Diagnostic and statistical manual of mental disorders (DSM-5). Here, we use ASD and autism interchangeably. Autism affects more than 5 million Americans, with an estimated prevalence of approximately 1 in 40 in children(Kogan et al., 2018), and is more commonly expressed in males(Bjørklund, Tinkov, et al., 2020; Pinto et al., 2010). The increasing number of affected individuals and families places a heavy burden on the healthcare system(Candon et al., 2018; Grosse et al., 2021). While the diagnosis of autism is well-defined, the underlying mechanisms are likely diverse and still poorly understood(Blatt & Fatemi, 2011; Nair et al., 2013, 2015; Puts et al., 2017), which hampers the development of treatments. A variety of genetic, environmental, and immunological factors likely contribute to the etiology of autism(Hallmayer et al., 2011; Ye et al., 2017).

In children, rapid cell differentiation associated with growth is linked to oxidative stress (OS). OS negatively affects cell function mostly through the release of highly reactive oxygen species (ROS) within the cell environment(Bjørklund, Meguid, et al., 2020; Chauhan et al., 2012). Free radicals are a natural, but undesirable, biproduct of cellular energy metabolism. Cells, therefore, employ antioxidants—the most abundant of which in the brain is glutathione (GSH)–to neutralize ROS by serving as electron donors to limit cell damage(Dringen, 2000; Dringen & Hirrlinger, 2003; Shimizu et al., 1998). Children’s developing brains are highly vulnerable to OS, due to greater immaturity of cells, higher oxygen consumption, and more unsaturated fatty acids(Ikonomidou & Kaindl, 2011). Increased OS arising from genetic and environmental factors may contribute to the neurophysiology of autism(Bjørklund, Meguid, et al., 2020; Chauhan et al., 2012; Janet K et al., 2017; Nadeem et al., 2019). Indeed, studies have shown that increased OS predicts reduced executive function and cognitive capacity(Frederic et al., 2015; Hajjar et al., 2018), reduced verbal fluency(Pia et al., 2016), reduced motor control(Hockenberry et al., 2015; Rodgers et al., 2016), and increased social anxiety(Bouayed et al., 2009), common in autism(Baron-Cohen & Belmonte, 2005; Miyajima et al., 2018; Mostofsky et al., 2007). Glutathione occurs in two forms to balance redox reactions and thereby to protect the cells from OS; a reduced form of GSH dimerizes to its oxidized form, glutathione disulfide GSSG(Owen & Butterfield, 2010), such that the ratio of GSH to GSSG gives information about the cells’ healthy response to OS. Both plasma(Geier et al., 2009; James et al., 2009; Nasrallah & Alzeer, 2022) and post-mortem studies(Chauhan et al., 2012; Rose et al., 2012) have shown lower GSH in autistic children compared to non-autistic children.

The majority of work investigating OS in autism has focused on peripheral markers, largely employing invasive and *in vitro* techniques. It is possible, however, to measure levels of endogenous GSH in the brain noninvasively using *in vivo* magnetic resonance spectroscopy (MRS). Preliminary MRS work(Durieux et al., 2016; Endres et al., 2017) has shown no GSH differences in autistic adults, but it is questionable whether their methodology can reliably resolve GSH signals which have low intensity and are heavily overlapped. J-difference-edited MRS(Harris et al., 2017; Mescher et al., 1998; Rothman et al., 1992) improves the resolution of GSH signals using frequency-selective pulses and subtraction. In the HERMES (Hadamard Encoding and Reconstructions of MEGA-edited Spectroscopy) experiment, a four-step editing scheme is used, effectively conducting two MEGA-PRESS experiments simultaneously. This editing scheme ensures that each target signal is treated independently, enabling the extraction of distinct edited spectra from the dataset in the different Hadamard combinations. HERMES has been applied to separate gamma-aminobutyric acid (GABA) and GSH(Maria et al., 2021; Oeltzschner et al., 2018; Saleh et al., 2016), as well as GABA, GSH and ethanol(Saleh, Wang, et al., 2020). GABA is also of keen interest in autism as the primary inhibitory neurotransmitter in the brain. Disturbed cortical excitation-inhibition balance is a persuasive hypothesis in autism(Hussman, 2001; Rubenstein & Merzenich, 2003) involving either or both upregulated glutamatergic excitatory and down-regulated GABAergic inhibitory systems(Sohal & Rubenstein, 2019). Multiple studies have found increased glutamate (Glu)(Brown et al., 2013; Hollestein et al., 2021, p.; Siegel-Ramsay et al., 2021) or reduced GABA(Gaetz et al., 2014; Puts et al., 2017; Rojas et al., 2014) in autism, though more recent reports failed to find Glu and GABA differences in autism(Brix et al., 2015; He et al., 2021; Kolodny et al., 2020). To date, no study has investigated GSH levels in autistic children using HERMES.

Differences in brain metabolite levels using MRS have been reported for autistic individuals in multiple brain regions including: the primary sensorimotor cortex SM1(He et al., 2021; Puts et al., 2017); the medial prefrontal cortex mPFC(Carvalho Pereira et al., 2018); the supplementary motor area SMA(Umesawa et al., 2020); the frontal lobe(Harada et al., 2011); auditory regions(Gaetz et al., 2014); the anterior cingulate cortex(Ito et al., 2017); and the thalamus(Friedman et al., 2003; He et al., 2021; Hegarty et al., 2018). All of these regions have been associated with autistic symptoms. For example, atypical metabolite concentrations in SM1, thalamus, mPFC, and SMA may relate to sensory and motor symptoms common to autism(Ford & Crewther, 2016; Margari et al., 2018). Nearly all incoming sensory information is filtered by the thalamus(Gomes et al., 2023; McCormick & Bal, 1994). SM1 plays crucial roles in the processing of sensory and motor information(Mostofsky et al., 2007; Puts et al., 2017), as well as motor control and learning(Ikeda et al., 2000). mPFC is a key region involved in executive function and adaptive behavior and appears important for aspects of social cognition, including theory of mind(Zhang et al., 2021), all commonly different in autism. SMA is a critical region for motor control and planning(Cona & Semenza, 2017; Mostofsky & Simmonds, 2008), through a number of thalamocortical circuits; and indeed atypical thalamocortical connectivity in autism has been reported previously(Green et al., 2017; Khan et al., 2016; Nair et al., 2013). Additionally, differences in volume and abnormal functional responses have been observed in autism across these cortical brain regions (SM1(Kleijer et al., 2018), mPFC(Courchesne et al., 2011; Kirkovski et al., 2016), SMA(Mostofsky et al., 2007, 2009)) and the thalamus(Hardan et al., 2006; Tamura et al., 2010).

Generally, the ^1^H-MRS studies show inconsistent findings due to different metabolites, age of the participants, the methodological techniques, and the brain region of interest. Here, we used the HERMES(Saleh, Wang, et al., 2020) MRS to measure both GSH and GABA levels in SM1, Thalamus (Thal), mPFC, and SMA in a pediatric cohort of autistic and non-autistic participants. We hypothesized that GSH and GABA concentrations would be significantly lower in autism compared to TDC in all four brain regions.

## 2. Methods

### 2.1 Participants

In all, this study recruited 101 child participants between 8-12 years old that were either autistic (ASD) or without neurodevelopmental conditions (TDC). The four brain regions were acquired in overlapping subsets of the whole cohort: SM1: 52 ASD (4F/48M) and 49 TDC (12F/37M); Thal: 17 ASD (1F/16M) and 16 TDC (1F/15M); mPFC: 22 ASD (2F/20M) and 21 TDC (6F/15M); SMA 27 ASD (3F/24M) and 29 TDC (10F/19M). Informed consent was obtained from a parent of each child (who also assented to study participation). All data were acquired with approval from the Johns Hopkins School of Medicine Institutional Review Board. Efficient methods were used to motivate participant compliance during scanning, including watching a movie, and getting a prize if they stayed still throughout the scan duration.

Children were excluded if they had a history of intellectual disability, brain injuries, seizures, known causes of autism (e.g. Fragile X), or other neurological disorders (e.g. Tourette syndrome). Autistic children with a co-occurring diagnosis of attention-deficit/hyperactivity disorder (ADHD, 32 ASD), anxiety disorders (1 ASD), oppositional defiant disorder (ODD, 4 ASD), dysthymia, or adjustment disorder (2 ASD) were not excluded. However, participants who had a history of, or met criteria for, conduct disorder, bipolar disorder, mania, or schizophrenia were excluded. Children taking stimulant medication temporarily ceased their stimulant medication the day before, and the day of, their MRI scan. Children included in the study met criteria for autism on both the Autism Diagnostic Observation Schedule-2 (ADOS-2)^111^ and the Kiddie Schedule for Affective Disorders and Schizophrenia (K-SADS)^110^. Children in the TDC group were excluded if they had a history of, or met criteria for, major depressive disorder, ADHD, ASD, anxiety disorders, or ODD. Children were also excluded from the TDC group if they had an immediate family member with ASD. Diagnostic information was reviewed and verified by an experienced child neurologist (SHM). See Table 1 for demographics.

**Table 1:**
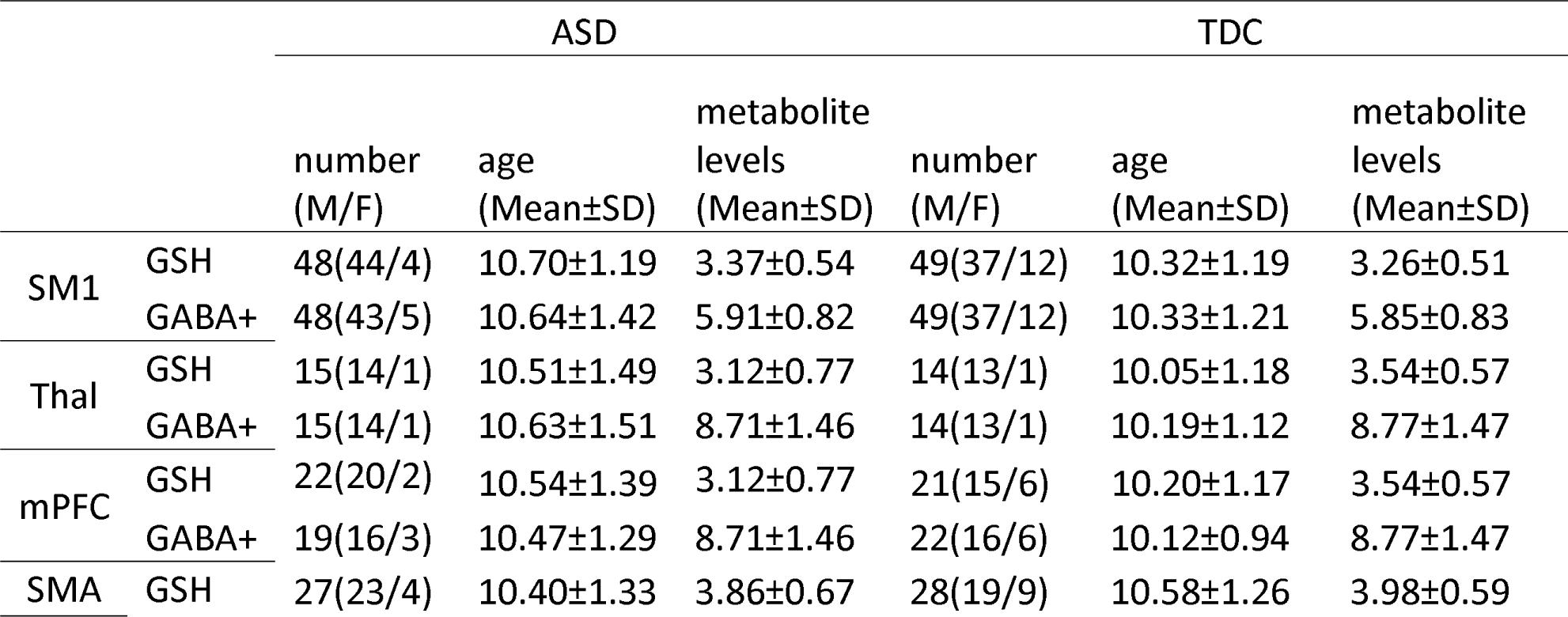
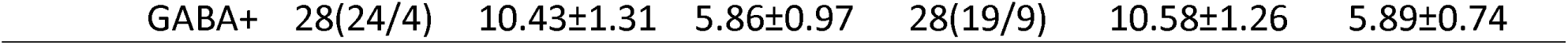
Mean and standard deviation (SD) GSH and GABA+ levels (in institutional units) in the four brain regions. GSH and GABA+ levels are presented for each brain region based on diagnosis group (ASD or TDC). Sex distribution and mean age are also listed for each brain region based on diagnosis group (as the number of participants collected varied for each brain region; see Methods for details). M=male; F=female.

Behavioral measures: The following standardized behavioral measures were completed by primary caregivers for both the ASD and TDC groups: the Adaptive Behavior Assessment System-II (ABAS-II)(*Harrison: Adaptive Behavior Assessment System - Google Scholar*, n.d.) or the updated version, the ABAS-3 (N=34), and The Social Responsiveness Scale (SRS-2)(*Constantino: Social Responsiveness Scale: SRS-2 - Google Scholar*, n.d.) groups. The ABAS-II and ABAS-3 are checklists used to access specific conceptual, social, and practical adaptive skills. The SRS-2 is a 65-item questionnaire that can be administered to a parent, which assesses a child’s ability to engage in emotionally appropriate reciprocal social interactions in naturalistic settings and includes items that ascertain social awareness, social information processing, capacity for reciprocal social responses, social anxiety/avoidance, and characteristic autistic preoccupations/traits. The SRS-2 generates a single score that can be used as a measure of severity of social function. Additionally, ADOS-2 measures were exclusively administered to the autism group. ADOS symptom clusters, including social interaction, communication, and stereotyped behaviors, were analyzed. Higher scores within these clusters were indicative of heightened symptom severity. Since many autistic children have a co-occurring diagnosis of ADHD, primary caregivers completed the Conners-3(K.C., n.d.-a) or Conners-4(K.C., n.d.-b) assessment to evaluate symptoms and impairments associated with ADHD using dimensional measures within this cohort. Scores corresponding to DSM-5 symptom levels were used to assess both Inattention and Hyperactivity/Impulsivity. Approximately 70% of the autistic children enrolled in our studies also exhibit comorbid ADHD, making this exploration particularly relevant.

### 2.2 Data acquisition

MRI: All data were collected on a Philips 3T Ingenia Elition RX scanner (Philips Healthcare, Best, the Netherlands) equipped with a 32-channel receive-only head coil. The scanning protocol included a 1 mm^3^ isotropic *T*_1_ MPRAGE for voxel placement and segmentation. MRS voxel sizes were: 30 (AP) x 30 (CC) x 30 (RL) mm^3^ in SM1, 26 (AP) x 24 (CC) x 40 (RL) mm^3^ in Thal, 30 (AP) x 20 (CC) x 40 (RL) mm^3^ in mPFC, and 30 (AP) x 30 (CC) x 30 (RL) mm^3^ in SMA, as shown in Figure 1. MR spectra were acquired using the following parameters: PRESS localization; *TR*/*TE*=2000/80 ms; 256 transients; HERMES editing with 20 ms editing pulses applied at 1.9 ppm and 4.56 ppm(Saleh, Wang, et al., 2020); 2048 datapoints; 2 kHz spectral width; and VAPOR water suppression(Tkáč et al., 1999) and interleaved water reference correction(Edden et al., 2016). The SM1 voxel was centered on the right central sulcus, posterior to the hand-knob(Yousry et al., 1997) in the axial plane and rotated to align with the cortical surface(Puts et al., 2011). The Thalamus voxel was placed parallel to the anterior commissure– posterior commissure line and covered the thalamus. The mPFC voxel was placed medially, anterior to the genu of the corpus callosum, and aligned with the anterior-to-posterior commissure line. The SMA voxel was placed symmetrically over the midline with its posterior face anterior to the central sulci(Boy et al., 2010).

**Figure 1:**
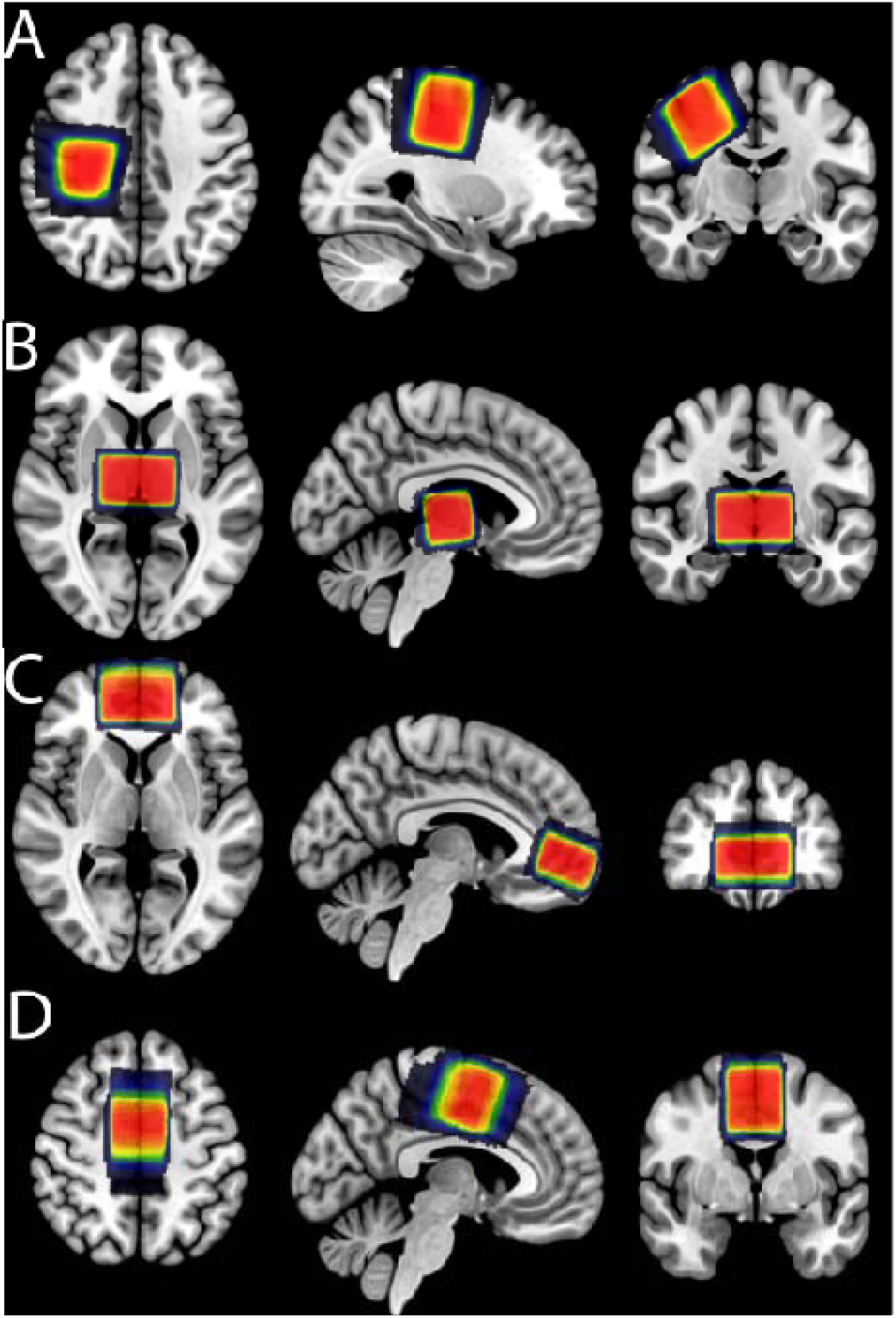
Location of the SM1 (A), Thal (B), mPFC (C), and SMA (D) voxels. Here, to illustrate anatomical overlap in voxel positioning between participants, each participant’s native space binary voxel mask was normalized to MNI space and overlaid onto the spm152 template (though metabolite concentrations used in statistical analyses were extracted from the voxel location in native space for each participant). Warmer colors (red color) indicate areas of greater overlap across participants.

### 2.3 Data analysis

All data were analyzed in Osprey software package (version 2.4.0)(Oeltzschner et al., 2020), using consensus-recommended pre-processing(Choi et al., 2021) and linear-combination modeling. HERMES data were frequency-and phase-corrected using probabilistic spectral alignment(Near et al., 2015). A Hankel singular value decomposition (HSVD) filter(Barkhuijsen et al., 1987) was applied to remove residual water signals and to reduce baseline roll. The modeled spectral range was from 0.2 to 4.2 ppm. A metabolite basis set was generated using MRSCloud(Hui et al., 2022), which included the following metabolites: ascorbate (Asc), aspartate (Asp), creatine (Cr), GABA, glycerophosphocholine (GPC), GSH, glutamine (Gln), Glu, myo-inositol (mI), lactate (Lac), N-acetyl aspartate (NAA), N-acetyl aspartyl glutamate (NAAG), phosphocholine (PCh), phosphorcreatine (PCr), phosphoryl ethanolamine (PE), scyllo-inositol (sI), and taurine (Tau). Eight macromolecule (MM) basis functions in the sum spectrum were included (MM_0.94_, MM_1.22_, MM_1.43_, MM_1.70_, MM_2.05_, Lip_0.9_, Lip_1.3_, Lip_2.0_). Co-edited MM peaks were defined at 1.2 and 1.4 ppm for the GSH-edited difference spectra, and at 0.93 and 3.0 ppm for the GABA-edited difference spectra. Amplitude-ratio soft constraints were applied to the amplitudes of the MM, lipids, and NAAG/NAA peaks as defined in the LCModel manual(Provencher, n.d.). The default Osprey baseline knot spacing of 0.4 ppm was used. Previous work(Choi et al., 2021; Craven et al., 2022; Song et al., 2022; Zöllner, Tapper, et al., 2022) has shown one-to-one amplitude-ratio soft constraint between the GABA amplitude and the co-edited macromolecules at 3.0 ppm (‘1to1GABAsoft’ in Osprey) performs better for edited MRS regarding to modeling performance. The water-reference data were quantified in the frequency domain with a six-parameter model (amplitude, zero-and first-order phase, Gaussian and Lorentzian line broadening, and frequency shift). A summary of the experimental methods and data quality, following the minimum reporting standards in MRS(Lin et al., 2021), is presented as Supplemental Table 1. All metabolite levels that were statistically analyzed were corrected for tissue composition. That is, for each participant, GSH levels were estimated after correcting for partial volume effects in each voxel using the method proposed in Gasparovic et al(Gasparovic et al., 2018), and GABA levels were alpha-corrected(Harris et al., 2015), since metabolite levels are reported to differ between brain gray and white matter(Gong et al., 2022). All reported GABA values in this study refer to GABA+, including the modeled co-edited macromolecules(Mullins et al., 2014).

### 2.4 Statistical analysis

Statistical analyses were conducted in RStudio (version 4.2.1; RStudio Team, 2021). GSH or GABA concentrations equal to 0 were interpreted as evidence of failure to fit and those data points were excluded from further analysis (SM1 n=0, Thal n=0, mPFC n=2, SMA n=2). The normality of metabolite distributions was assessed using the Shapiro-Wilk test. In order to conduct a comparative analysis of the ABAS and SRS-2, a one-way Analysis of Variance (ANOVA) was employed. Due to the observed disparity in male-to-female ratios between the two groups, we decided to employ the robust linear mixed-effects model (LME). Two robust regression LME models (package(nlme)(Pinheiro et al., 2023)) were estimated for each brain region to investigate the relationship between metabolite (GSH or GABA+) levels and diagnosis group, age, and sex, accounting for the random intercept of each participant. Pearson correlations were also calculated between metabolite levels and dimensional scores of ASD and ADHD symptom severity (respectively Conners and ADOS for the ASD group only). We considered statistical significance as *p*<0.05.

## 3. Results

Metabolite levels for the four examined brain regions are summarized in Table 1, including the mean age and sex distribution for each sample for each brain region. Some spectra were excluded due to out-of-voxel echoes in the spectra(Ernst & Chang, 1996; Song et al., 2022). Specifically, three GSH-edited (2 ASD and 1 TDC) and three GABA-edited spectra (2 ASD and 1 TDC) were excluded from the SM1 region, four GSH-edited and (2 ASD and 2 TDC) four GABA-edited spectra (2 ASD and 2 TDC) from the Thal region, ten GSH-edited (5 ASD and 5 TDC) and twelve GABA-edited spectra (8 ASD and 4 TDC) from the mPFC region, and one GSH-edited (1 ASD) spectrum from the SMA region. After these exclusions, we included in subsequent statistical analyses SM1: 97 GSH and 97 GABA+, Thal: 29 GSH and 29 GABA+, mPFC: 43 GSH and 41 GABA+, and SMA: 55 GSH and 56 GABA+. See Table 1 for demographics.

GSH-and GABA-edited HERMES difference spectra, are overlaid for all participants in each group in Figure 2, providing a comprehensive visualization of the metabolic profiles under investigation. The GSH peak at 2.95 ppm and the GABA+ peak at 3.0 ppm are prominently displayed, representing the key markers of interest in our analysis. The black line signifies the mean spectra within each group, as a central reference point when considering the individual subject spectra. Figure 3 shows boxplots of GSH and GABA+ levels by diagnosis group for the four brain regions considered in this study. LME results are displayed in Table 2. There were no statistically significant associations between GSH or GABA+ levels and diagnosis group, age, or sex within any of the four examined brain regions (SM1, Thal, mPFC, or SMA); *p*>0.05 in all cases. Note, we reported a subset of the SM1 GABA+ data (from 24 ASD and 26 TDC participants) and Thal GABA+ data (from 16 ASD and 9 TDC participants) in our previous work(He et al., 2021).

**Figure 2:**
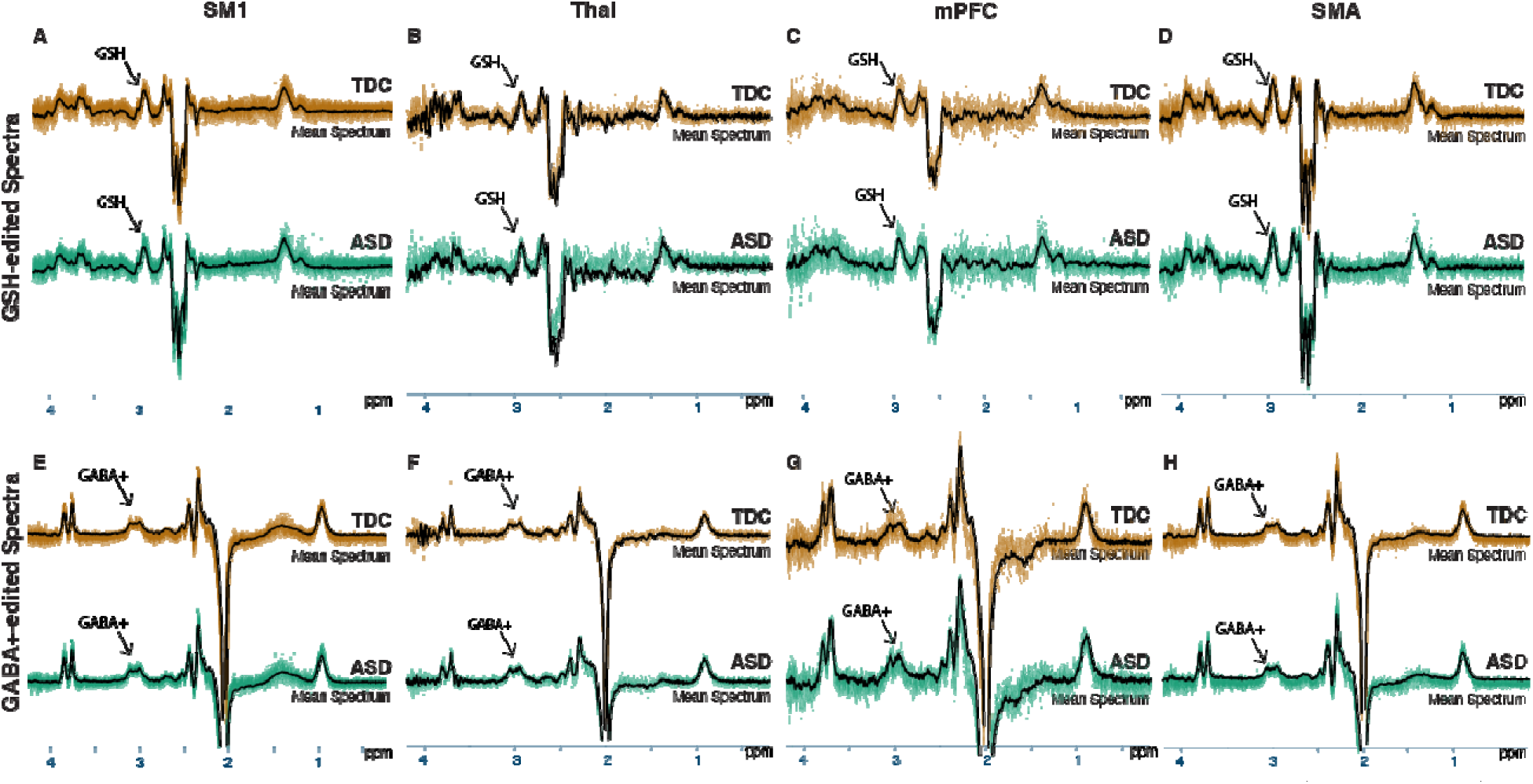
*In vivo* GSH-edited and GABA+-edited spectra acquired from the SM1 (A, E, n=97), Thal (B, F, n=29), mPFC (C, G, n=43(GSH)/41(GABA)), and SMA (D, H, n=55(GSH)/56(GABA)) regions. The top row (A–D) shows the GSH-edited spectra and the bottom row (E–H) shows the GABA+-edited spectra. The black line within each spectrum represents the mean spectrum of each group. The orange spectra are those acquired from the TDC group, while the green spectra are those acquired from the ASD group. Arrows point to the edited GSH and GABA+ peaks within the respective spectra.

**Figure 3:**
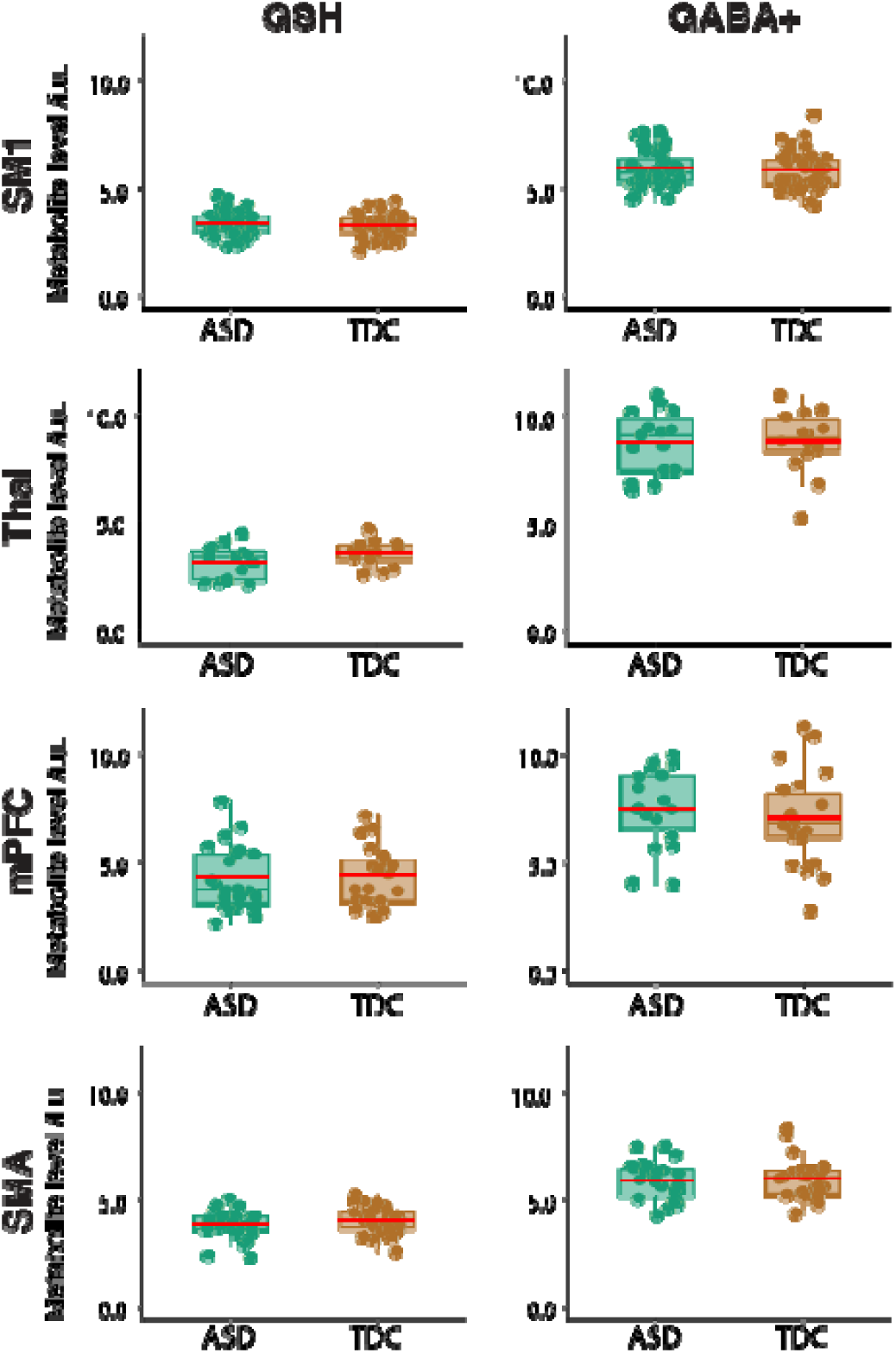
Boxplots of GSH (left panel) and GABA+ levels (right panel) in the SM1, Thal, mPFC, and SMA regions for the ASD (green) and TDC (orange) groups. The red line in each box represents mean metabolite levels.

**Table 2:**
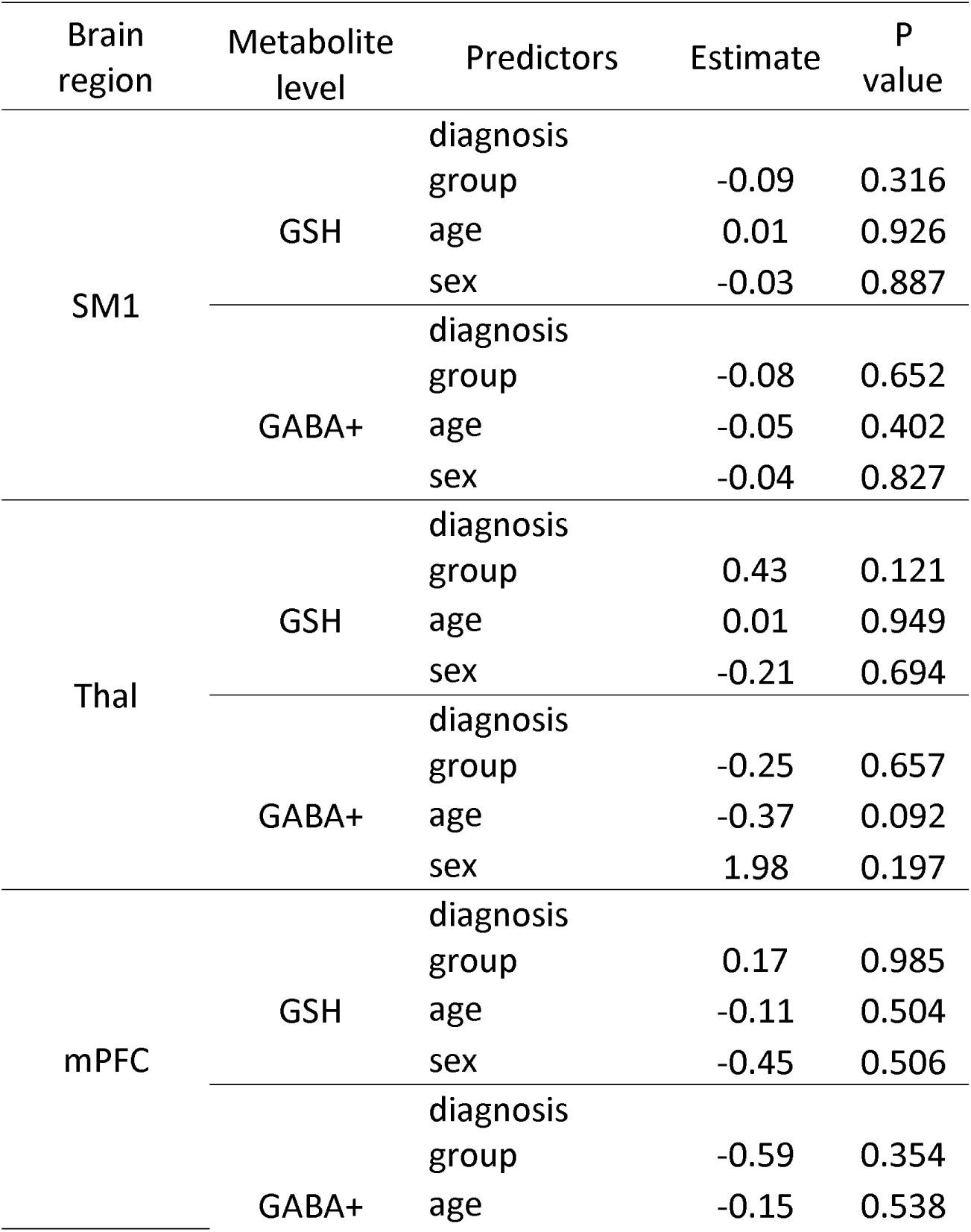
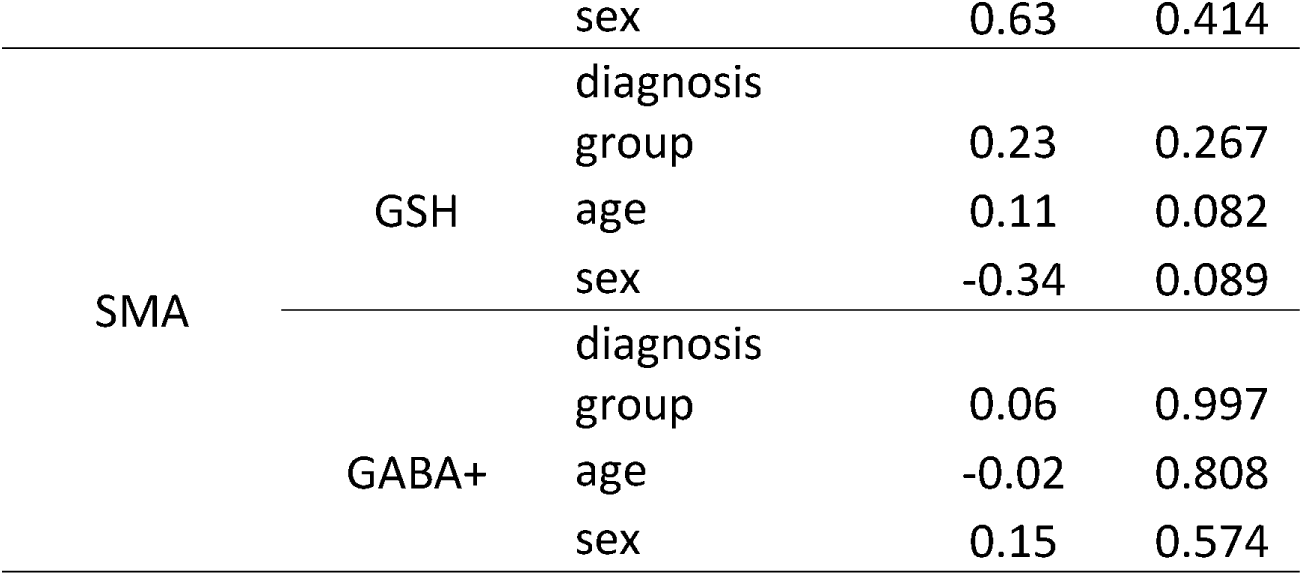
Variation of regional brain metabolite levels with diagnosis group, age, and sex as fixed effects in robust linear mixed-effects (LME) models. There were no relationships between autism diagnosis, age, or sex and GSH or GABA+ levels for any brain region (*p*>0.05 in all cases).

As anticipated, compared to the TDC group, the ASD group showed overall lower scores (indicative of worse symptoms) in adaptive behavior ratings (Global Adaptive Composite, GAC scores) (p<0.001) and showed higher SRS-P scores (shown in Supplemental Table 2). There were no significant correlations between GSH or GABA+ levels in any of the four brain regions and autism symptomology or behavior based on the ADOS-2 or SRS-P or ABAS assessments (Correlation: 0.01-0.25(ADOS-2), 0-0.2 (SRS-P), and -0.5-0.2(ABAS), all p>0.05); See Supplemental Figure 1&2. Higher GABA levels in SM1 region (correlation coefficient=0.52; p<0.001) and Thal region (correlation coefficient=0.56; p=0.01) were significantly correlated with higher inattention scores from the Conners rating scale, in spite of the fact that there is no correlation between GABA levels in the two regions, while other regions were not significant significantly correlated. There were no significant correlations between metabolite levels and Conners hyperactivity/impulsivity scores (Correlation: -0.15-0.19, all p>0.05). See Supplemental Figure 3.

## 4. Discussion

This is the first study to simultaneously quantify both cerebral GSH and GABA+ levels *in vivo* in a large sample of autistic children using HERMES edited spectroscopy at 3T. Contrary to our hypotheses, we observed no significant group differences in either GSH or GABA+ concentrations in any of the four brain regions of interest: SM1, Thal, mPFC, and SMA. Our GSH findings are consistent with previous MRS studies reporting no GSH differences in adults with autism in subcortical(Endres et al., 2017) or cortical brain regions(Durieux et al., 2016; Endres et al., 2017), although both of these studies measured GSH using a short-*TE* STEAM acquisition rather than our more specific spectral editing approach. Edited MRS (i.e. Hadamard Encoding and Reconstructions of MEGA-edited Spectroscopy HERMES, which was used in this study), allows us to differentiate GSH from other overlapping signals, providing better specificity and accuracy of the measured metabolite levels, and thus increased reliability of our results. In the HERMES experiment, a four-step editing scheme is used, effectively conducting two MEGA-PRESS experiments simultaneously. This editing scheme ensures that each target signal is treated independently, enabling the extraction of distinct edited spectra from the dataset in the different Hadamard combinations. Hadamard-encoded editing offers the advantage of simultaneous editing for two or more metabolite targets, which has been applied to separate GABA and GSH(Maria et al., 2021; Oeltzschner et al., 2018; Saleh et al., 2016), as well as GABA, GSH and ethanol(Saleh, Wang, et al., 2020). HERMES addresses one significant limitation of edited MRS – the extended scan times required to make multiple single-metabolite measurements.

The study conducted by Endres et al.(Endres et al., 2017) examined a left dorsolateral prefrontal cortex DLPFC voxel of 15 mL and a dorsal anterior cingulate cortex voxel of 15.6 mL in 24 adults with ASD (M/F 12/12) and 18 healthy controls (M/F 8/10). Durieux et al.(Durieux et al., 2016), on the other hand, explored basal ganglia (6 mL) and DLPFC (12.8 mL) regions in 21 men with ASD and 29 healthy male controls. In our study, we opted for larger voxels (24-27 mL) to improve the accuracy of quantifying low-concentration metabolites (i.e. GSH and GABA+) in the developing brain. None of these studies revealed any significant differences between the ASD group and the healthy control group, although it is appropriate to be cautious in interpreting negative findings. GSH has limited permeability through the blood-brain barrier(Haddad et al., 2021), a systemic isolation which might contribute to the stability of cerebral GSH levels with pathology(Haddad et al., 2021) and aging(Gong et al., 2022; Saleh, Papantoni, et al., 2020; Tong et al., 2016). Consistent with our findings, post-mortem measurements in a cohort of twenty individuals (10 ASD and 10 TDC, aged 5-38 years old) also did not show significant differences in GSH levels in the frontal, parietal, or occipital cortices(Chauhan et al., 2012), a useful validation of challenging *in vivo* MRS measures. The lack of clear GSH differences suggests that alterations in cerebral GSH metabolism may not be a prominent factor in the pathophysiology of autism (or at least, not one that persists throughout life), even though previous research(Bjørklund, Meguid, et al., 2020; Chauhan et al., 2012; Janet K et al., 2017; Nadeem et al., 2019) has proposed a link between OS and autism, with GSH being a crucial antioxidant in the brain.

Most GSH studies in the literature employ *in vitro* blood assays, while our study focused on cerebral GSH levels, employing non-invasive edited MRS. Although our cerebral GSH results do not support the redox hypothesis of autism in the brain, previous results of decreased GSH in the plasma of autistic children may help explain some other inflammatory symptoms that autistic children experience (e.g. gastrointestinal discomfort(Taniya et al., 2022)). For instance, one previous study found decreased GSH in plasma in autistic children(Nasrallah & Alzeer, 2022), providing some evidence for the redox hypothesis in autism. Recent clinical trials(Hardan et al., 2012; Hendren et al., 2016) support a possible redox imbalance in autistic children. Autistic symptoms (i.e. an irritability subscale of the Aberrant Behavior Checklist (ABC)(Norris et al., 2019)) improved significantly following increased levels of cysteine (a GSH precursor), leading to an increased GSH plasma concentration(Hardan et al., 2012). Unlike GSH, N-acetylcysteine NAC crosses the blood-brain barrier and might be expected to influence systemic and cerebral GSH levels. A future study should compare levels of GSH in plasma and brain following NAC administration.

Our findings of no significant GABA+ differences between autistic children and TDC are consistent with several recent *in vivo* studies comparing GABA+ levels between autistic and TDC groups in the thalamus(He et al., 2021), ACC(Ajram et al., 2017; Cochran et al., 2015; Goji et al., 2017), frontal cortex(Ajram et al., 2017; Brix et al., 2015), SM1(He et al., 2021; Robertson et al., 2016), left lenticular nucleus(Harada et al., 2011), occipital cortex(Drenthen et al., 2016; Gaetz et al., 2014; Puts et al., 2017; Robertson et al., 2016, 2016), and cerebellum(Goji et al., 2017), at 3T using the MEGA-PRESS sequence. However, findings from the literature related to GABA+ levels in autism are mixed; some studies (including our own) have reported lower GABA+ levels in ACC (Ito et al., 2017), frontal cortex(Harada et al., 2011), SM1(Gaetz et al., 2014; Puts et al., 2017), temporal cortex(Gaetz et al., 2014; Port et al., 2017; Rojas et al., 2014), and cerebellum (Ito et al., 2017) in autistic children using MEGA-PRESS at 3T. This, in fact, is in line with post-mortem studies(Palmen et al., 2004; Williams et al., 1980) reporting decreased numbers of GABAergic Purkinje cells in the cerebellar cortex of children with autism. One MRS study(Maier et al., 2022) and a number of plasma-based studies(Alabdali et al., 2014; Al-Otaish et al., 2018; Dhossche et al., 2002; Russo, 2013; Saha et al., 2022) found elevated GABA levels in autistic children compared to TDC; these studies also found positive correlations between plasma GABA levels and severity of autistic symptoms, including levels of hyperactivity and tactile sensitivity(Alabdali et al., 2014; Russo, 2013). The inconsistency across GABA studies could be attributed to the complexity and heterogeneity of autism itself, different brain regions studied (as well as differing approaches to delineating brain regions), varied participant ages, and methodological heterogeneity.

While it is perhaps somewhat surprising that no group differences in metabolite concentrations were found, either for GABA (as has been shown before) or for GSH, we note that autism is an extremely heterogeneous condition, and that group differences, if present, may be small and relatively uninformative in understanding the broader autism phenotype(Loth et al., 2021). Subgroup approaches may be better suited to determine whether there are indeed individuals with specific neurometabolic differences, and whether these are subsequently associated with differences in clinical outcomes and lived experiences. The only significant relationship that was observed between behavioral symptom assessments and metabolite levels was the positive correlation between GABA levels, both in SM1 (correlation coefficient=0.52, p<0.01) and Thal regions (correlation coefficient=0.52, p<0.01), and ADHD inattention scores. This finding aligns with a task-dependent study(Mamiya et al., 2022) that indicates a potential connection between attention control deficits in ADHD and disorder in the GABAergic system within the brain.

One limitation in this study was that some participants continued treatment of psychotropic medication (e.g., fluoxetine, atomoxetine, etc.) during the study. These medicines are reported to have no significant effect on GSH and GABA+ levels(Puts et al., 2017); thus, for both ethical and practical reasons, participants did not discontinue these medications before participation in the study (though they did discontinue stimulant medications the day prior to and day of the study; see Methods for details). Another limitation of the present cohort is the under-representation of autistic girls. The male-to-female ratio is 4:1 in autistic populations without intellectual impairment(Loomes et al., 2017; Mandy et al., 2012), while the cohort in our study has a male-to-female ratio around 10:1. Diagnostic tools for autistic populations are standardized on majority-male samples, which may lead to missed diagnoses in females^113^. Future studies should include a male-to-female ratio more representative of the autism population. The heterogeneity of our sample might be addressed in future studies with larger sample sizes to allow for comparison between more heterogeneous subgroups (in terms of medical history, comorbidities, and treatment regimen). Finally, one limitation of edited MRS is that the motion-sensitive acquisition from each region is carried out sequentially, resulting in long protocol durations. Some patients are not able to remain compliant throughout the protocol, which could lead to motion-related artifacts in the MRS data, and the potential for motion-driven false positive results. However, we visually inspected all spectra for the expected GSH and GABA+ peaks and conservatively excluded any spectra which contained visible artifacts such as OOVs (see Methods); further, all included spectra met consensus-recommended(Wilson et al., 2019) data quality thresholds (e.g., 12 Hz linewidth). The Cr frequency stability metric indicates that the standard deviation (SD) values for all included spectra are less than 0.2, with 95% of them being less than 0.1 (as shown in supplemental Table 3). This underscores the exceptional stability observed.

## Conclusion

Our study measured GSH and GABA+ levels in four brain regions in a large cohort of autistic and typically developing children. We did not find significant between-group differences in either metabolite in any brain region. These findings contribute to the growing body of knowledge on the neurochemical profiles associated with autism and provide valuable insights into the complex interplay of neurotransmitters and neurochemical signaling. While our results do not support the hypothesis of differential GSH and GABA+ levels in these regions between autistic and TDC individuals, it is important to note that further research is warranted to explore other potential neurochemical alterations and their relationship to the pathophysiology of autism.

## Acknowledgements

This work was supported by funding from the National Institute of Health (NIH) grants R00 AG062230, R01 EB016089, R01 EB023963, P41 EB031771, K99 AG080084, R21 AG060245, K99 AG080084, P50 HD103538(IDDRC), NSF 2124276.

