## Supplemental Table 1 for "Brain Glutathione and GABA+ levels in autistic children"

| | |
|---|---|
| 1.Hardware | |
| a. Field strength [T] | 3 T |
| b. Manufacturer | Philips |
| c. Model (software version if available) | R 5.7.1 |
| d. RF coils: nuclei (transmit/receive), number of channels, type, body part | $^{1}$H, 32 channel, head |
| e. Additional hardware | - |
| 2. Acquisition | |
| a. Pulse sequence | HERMES (Johns Hopkins University Patch) |
| b. Volume of interest (VOI) locations | SM1, Thal, mPFC, SMA |
| c. Nominal VOI size [mm$^3$] | SM1/ mPFC/ SMA:30 x 30 x 30 mm$^3$, Thal: 26(AP) x 24(CC) x 40(LR) mm$^3$ |
| d. Repetition time (TR), echo time (TE)[ms] | TR 2000 ms, TE 80 ms |
| e. Total number of averages per spectrum<br>i. Number of averaged specra per subspectrum | 320 total averages with 80 averages per subspectrum |
| f. Additional sequence parameters<br>i. Editing pulse parameters | F1: 2000 Hz, 2048 points<br>GABA at 1.9 ppm, GSH at 4.56 ppm |
| g. Water suppression method | VAPOR |
| h. Shimming method, reference peak, and threshold of acceptance of shim chosen | 1st and 2nd shimming pencil beam, water |
| i. Trigger or motion correction | No trigger or active motion correction |
| 3. Data analysis methods and outputs | |
| a. Analysis software | Osprey 2.4.0 |
| b. Processing steps deviating from Osprey | Final alignment of the averaged sub-spectra by minimizing the choline (not water) peak; co-edited MMs at 3 ppm were modelled using the “1to1GABAsoft” model |
| c. Output measure | tCr, rawWaterScaled, CSFWaterScaled, TissCorrWaterScaled |
| d. Quantification references and assumptions, fitting model assumptions | Asc, Asp, Cr, GABA, GPC, GSH, Gln, Glu, mI, Lac, NAA, NAAG, PCh, PCr, PE, scyllo-inositol, Tau, 8 MM basis functions in the sum spectrum ($MM_{0.94}$, $MM_{1.22}$, $MM_{1.43}$, $MM_{1.70}$, $MM_{2.05}$, Lip09, Lip13, Lip20)<br>Fitting method: Osprey baseline knot spacing 0.4 ppm |

| 4. Data quality | |
|---|---|
| HERMES, SM1 | |
| a. SNR (Cr), linewidth (Cr) [Hz, OFF spectra] | SNR 230 ± 41, linewidth 4.99 ± 0.78 Hz |
| c. Quality measures of postporcessing model fitting (Mean Relative Amplitude Residual (Residual/Noise))<br>sum<br>diff1<br>diff2 | <br><br><br>22.35 ± 16.98<br>2.59 ± 0.72<br>2.28 ± 0.90 |
| d. Mean spectrum created with OspreyOverview | Figure 2 |
| HERMES, Thal | |
| a. SNR (Cr), linewidth (Cr) [Hz, OFF spectra] | SNR 123 ± 31, linewidth 6.06 ± 0.79 Hz |
| c. Quality measures of postporcessing model fitting (Mean Relative Amplitude Residual (Residual/Noise))<br>sum<br>diff1<br>diff2 | <br><br><br>8.76 ± 4.69<br>4.51 ± 4.28<br>4.66 ± 3.67 |
| d. Mean spectrum created with OspreyOverview | Figure 2 |
| HERMES, mPFC | |
| a. SNR (Cr), linewidth (Cr) [Hz, OFF spectra] | SNR 106 ±29, linewidth 7.52 ± 1.54 Hz |
| c. Quality measures of postporcessing model fitting (Mean Relative Amplitude Residual (Residual/Noise))<br>sum<br>diff1<br>diff2 | <br><br><br>7.99 ± 6029<br>2.71 ± 0.57<br>2.84 ± 1.01 |
| d. Mean spectrum created with OspreyOverview | Figure 2 |
| HERMES, SMA | |
| a. SNR (Cr), linewidth (Cr) [Hz, OFF spectra] | SNR 220 ± 37, linewidth 4.67 ± 1.02 Hz |
| c. Quality measures of postporcessing model fitting (Mean Relative Amplitude Residual (Residual/Noise))<br>sum<br>diff1 | <br><br><br>16.05 ± 6.55<br>2.74 ± 0.63<br>2.29 ± 0.49 |

| | |
|---|---|
| diff2 | |
| d. Mean spectrum created with OspreyOverview | Figure 2 |

Supplementary Material 1: Summary following minimum reporting standards in MRS generated in Osprey See Lin et al. 'Minimum Reporting Standards for in vivo Magnetic Resonance Spectroscopy (MRSinMRS): Experts' consensus recommendations. NMR in Biomedicine. 2021;e4484. [doi.org/10.1002/nbm.4448](https://doi.org/10.1002/nbm.4448)
