## Supplemental Table 2 for "Brain Glutathione and GABA+ levels in autistic children"

| | ASD | | TDC | | Statistic |
|---|---|---|---|---|---|
| | Mean | SD | Mean | SD | |
| **ABAS[a]** | | | | | |
| Global Adaptive Composite | 74.36 | 12.83 | 106.03 | 9.73 | p< 0.001 |
| **SRS-P-2[aa]** | | | | | |
| SRS_P_2_Total_T_Score | 73.89 | 9.46 | 45.84 | 7.69 | p< 0.001 |
| **ADOS-2[aaa]** | | | | | |
| ADOS2_Total | 12.32 | 4.42 | | | |
| ADOS2_Comparison_Score | 7.06 | 1.82 | | | |
| ADOS2_Social_Affect_Total | 9.43 | 3.49 | | | |
| ADOS2_RRB_restricted_and_repetitive_behaviors_Total | 3.17 | 2.62 | | | |

ª n_ASD=44; n_TDC=39; ªª n_ASD=44; n_TDC=37; ªªª n_ASD=47
