## Supplementary figures and images for "Brain Glutathione and GABA+ levels in autistic children"

### Supplemental Figure 1

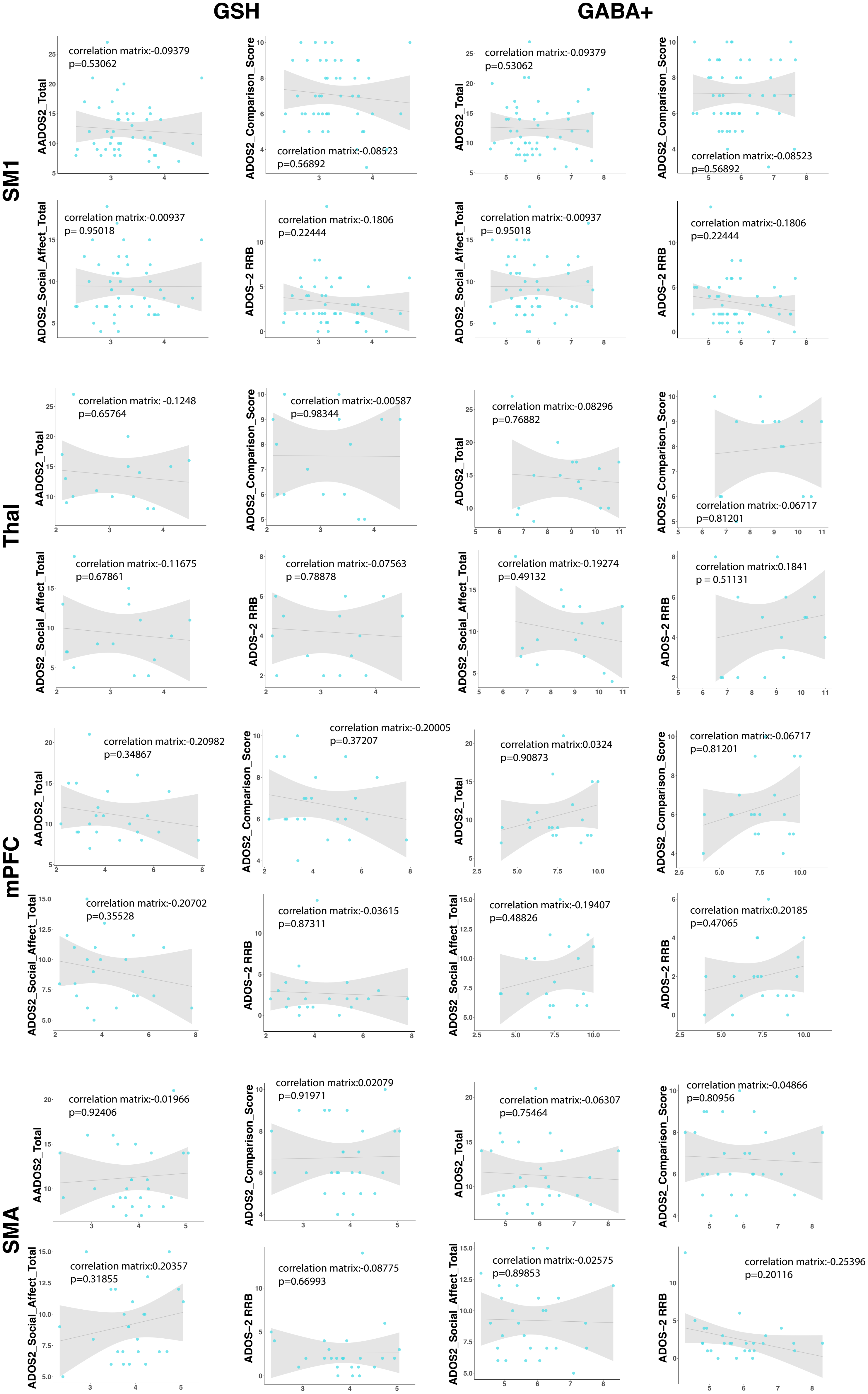

### Supplemental Figure 2

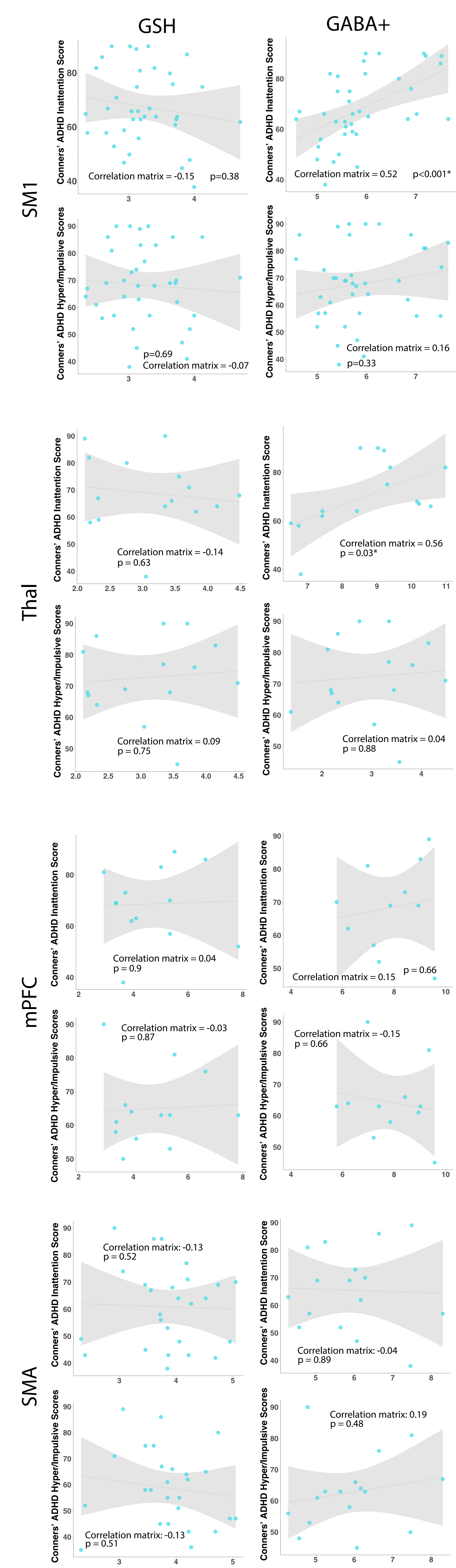

### Supplemental Figure 3

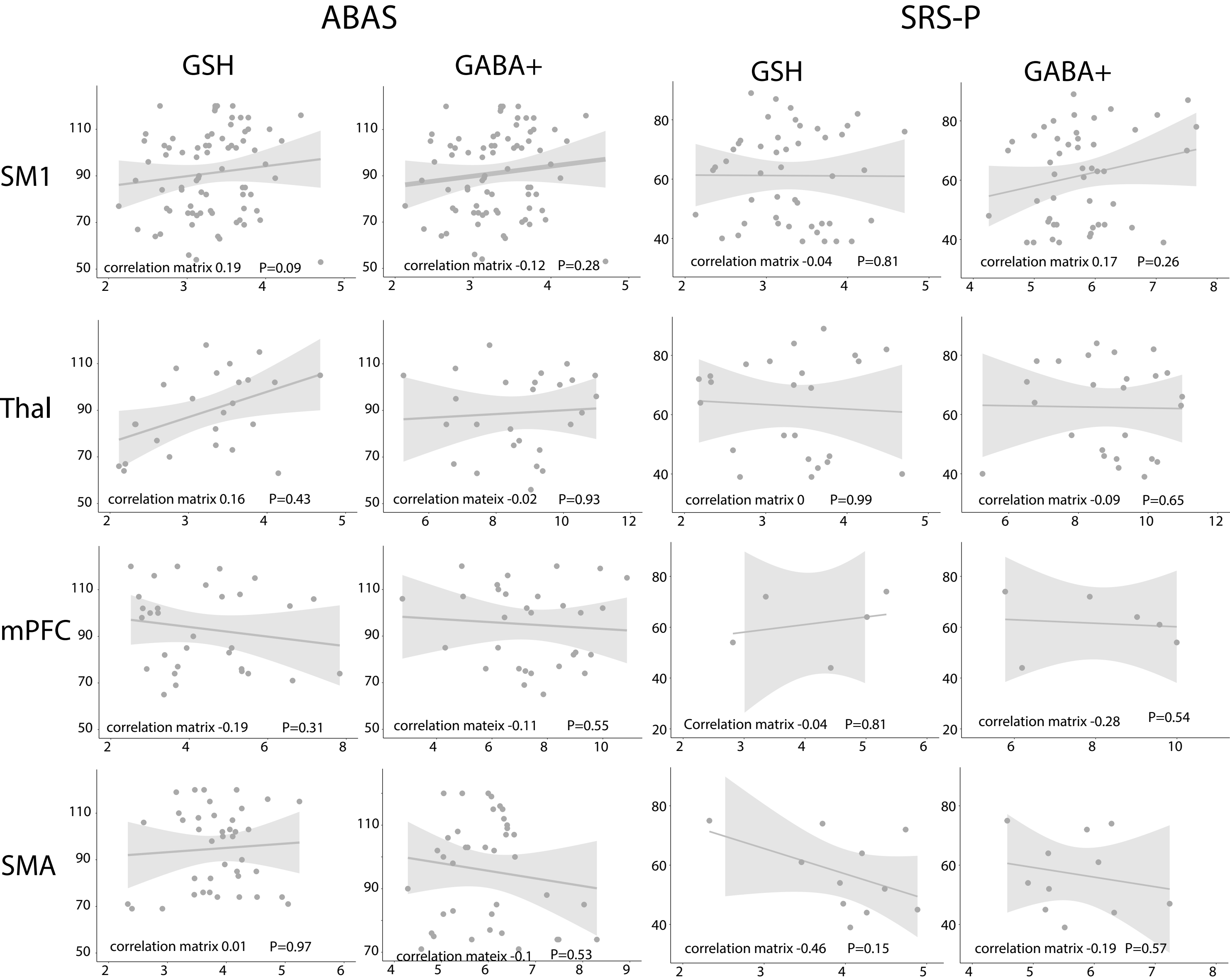
